# Sequential Optimal Experimental Design of Perturbation Screens Guided by Multi-modal Priors

**DOI:** 10.1101/2023.12.12.571389

**Authors:** Kexin Huang, Romain Lopez, Jan-Christian Hütter, Takamasa Kudo, Antonio Rios, Aviv Regev

## Abstract

Understanding a cell’s expression response to genetic perturbations helps to address important challenges in biology and medicine, including the function of gene circuits, discovery of therapeutic targets and cell reprogramming and engineering. In recent years, Perturb-seq, pooled genetic screens with single cell RNA-seq (scRNA-seq) readouts, has emerged as a common method to collect such data. However, irrespective of technological advances, because combinations of gene perturbations can have unpredictable, non-additive effects, the number of experimental configurations far exceeds experimental capacity, and for certain cases, the number of available cells. While recent machine learning models, trained on existing Perturb-seq data sets, can predict perturbation outcomes with some degree of accuracy, they are currently limited by sub-optimal training set selection and the small number of cell contexts of training data, leading to poor predictions for unexplored parts of perturbation space. As biologists deploy Perturb-seq across diverse biological systems, there is an enormous need for algorithms to guide iterative experiments while exploring the large space of possible perturbations and their combinations. Here, we propose a sequential approach for designing Perturb-seq experiments that uses the model to strategically select the most informative perturbations at each step for subsequent experiments. This enables a significantly more efficient exploration of the perturbation space, while predicting the effect of the rest of the unseen perturbations with high-fidelity. Analysis of a previous large-scale Perturb-seq experiment reveals that our setting is severely restricted by the number of examples and rounds, falling into a non-conventional active learning regime called “active learning on a budget”. Motivated by this insight, we develop IterPert, a novel active learning method that exploits rich and multi-modal prior knowledge in order to efficiently guide the selection of subsequent perturbations. Using prior knowledge for this task is novel, and crucial for successful active learning on a budget. We validate IterPert using insilico benchmarking of active learning, constructed from a large-scale CRISPRi Perturb-seq data set. We find that IterPert outperforms other active learning strategies by reaching comparable accuracy at only a third of the number of perturbations profiled as the next best method. Overall, our results demonstrate the potential of sequentially designing perturbation screens through IterPert.

## 1 Introduction

The expression response of a cell to a genetic perturbation reveals fundamental insights into cell and gene function [1]. Perturb-seq is a relatively recent technology for pooled genetic screens with a single-cell RNA seq (scRNA-seq) readout of the expression response to a perturbation [2, 3, 4]. Perturb-seq provides insights into gene regulatory machinery [5], helps identify target genes for therapeutic intervention [6], and can facilitate the engineering cells with a specific target state [7, 8]. Recent technical advances have enhanced the scope, scale and efficiency of Perturb-Seq [9, 4, 10]. However, because of the plethora of biological contexts, across cell types, states and stimuli, and the need to test combinations of perturbations (due to the possibility of non-additive genetic interactions), the number of required experiments explodes combinatorially. With trillions of potential experimental configurations or more, it becomes unrealistic to conduct all of them directly [8, 11].

Recently, researchers proposed machine learning models to predict perturbation outcomes [12, 13, 14]. Such models are trained on existing Perturb-seq datasets [10, 2, 8, 11] and then predict expression outcomes of unseen perturbations, of single genes or their combinations. While promising, these models suffer from a selection bias caused by the design of the original experiment used for training, in terms of selected perturbations and biological conditions. In particular, the training data are often profiled to answer a specific biological question, but not to maximize the predictive accuracy of the machine learning model across a large pool of unprofiled perturbations.

In this work, we present a novel paradigm for exploring a perturbation space by executing a sequence of Perturb-seq experiments. At the core of this paradigm lies a sequential optimal design procedure that inter-leaves the machine learning model and the wet-lab, where the Perturb-seq assay is performed. At each step of the sequence, we acquire data and use it to re-train the machine learning model. Then, we apply an optimal design strategy to select a batch of perturbation experiments that will most benefit the model to predict all of the unprofiled perturbations. The key idea is to sample the perturbation space intelligently by considering perturbations that are most informative and representative to the model, while accounting for diversity. Using this strategy, we can run as few perturbation experiments as possible, while obtaining a model that has sufficiently explored the perturbation space.

This idea is well-studied in the machine learning literature, and is the topic of active learning [15]. Active learning has been used in practice across many domains, such as document classification [16], medical imaging [17] and speech recognition [18]. However, we noticed that effective active learning approaches necessitate a substantial initial set of labeled examples (i.e., in our case, profiled perturbations), complemented by numerous batches that collectively result in tens of thousands of labeled data points [19, 20]. In contrast, the constraints of iterative Perturb-seq in the lab make such conditions unattainable, both in terms of cost and time (as shown in our economic analysis in Section 3.1). In this “budgeted” regime, it has been reported that random selection outperforms most active learning strategies [21, 22, 23].

We therefore propose a new strategy called IterPert (ITERative PERTurb-seq) that tackles the active learning on a budget setting for Perturb-seq data. Motivated by a data-driven analysis, our key observation is that when on a budget, it may be beneficial to combine the evidence from the data with publicly available sources of prior-knowledge, especially in the first few rounds. Such examples of prior knowledge include Perturb-seq data from related systems, large scale genetic screens with other modalities, such as genome-scale optical pooled screens [24, 25], and data on physical molecular interactions, such as protein complexes [26, 27]. This prior information spans multiple modalities such as networks, text, image, and 3D structure, which may be challenging to exploit during active learning. We overcome this by defining reproducing kernel Hilbert spaces on each of the modalities, and applying a kernel fusion strategy [28] to combine information from multiple sources.

To compare IterPert against other commonly used methods, we conducted an extensive empirical study using a large-scale single-gene CRISPRi Perturb-seq dataset collected in a cancer cell line (K562 cells) [11] and benchmarked 8 recent active learning strategies. IterPert achieved similar accuracy as the best active learning strategy but with three times fewer perturbations profiled as the training data. IterPert also showed robust performance in both essential genes screens and genome-scale screens, and when considering batch effects across iterations.

To summarize, our contributions are (1) proposing a sequential experimental design approach to Perturb-seq profiling for efficient exploration of a perturbation space; (2) identifying the algorithmic problem of active learning on a budget in this setting; (3) proposing a new active learning strategy that incorporates prior information and obtains a speedup of more than three times over the best baseline strategy.

## 2 Background

### Perturb-seq prediction model

We consider a predictive model *f*_*θ*_ with parameters *θ* that maps a set of perturbations 𝒫 = (*P*_1_, …, *P*_*M*_) to the post-perturbed expression outcome ŷ ∈ *ℝ*^*L*^, where *L* denotes the number of genes with measured expression levels. We denote the set of available Perturb-seq training data as 𝒟_train_ = 𝒳_train_ × 𝒴_train_, where 𝒳_train_ = 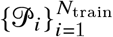 and 𝒴_train_ = 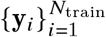, respectively.

Several models have been designed for this specific task [12, 13, 29, 30], and our proposed framework can be adapted for any of those (refer to Section 5). However, in the remainder of this paper we focus on adopting the current state-of-the-art model GEARS [12] as the prediction model for active learning. GEARS is a deep learning model customized for perturbation prediction that uses graph neural networks (GNN) to incorporate gene ontology and gene co-expression graphs to learn perturbation embeddings from data. GEARS uses a focal loss as the objective function during training in order to assign higher weight to differentially expressed genes: 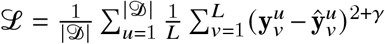, where *γ* = 2 and 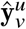, 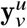 are the predicted and true expression level of gene *ν* after perturbation *u*, respectively.

### Batch-mode pool-based active learning

Except for the specific low-budget setting, the active learning problem we are interested in has been well-studied in the literature [15] and corresponds to batch-mode pool-based active learning. It can be formulated as follows: We consider an initial labeled training set 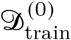 and an unlabeled pool set 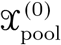. In each subsequent round *i*, we first train a model *f*_*θ*_ on 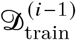. Then, an active learning selection strategy *g* takes in (1) a pre-specified batch size *N*_batch_, (2) the training set 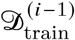, (3) the unlabeled pool set 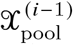, and (4) the model *f*_*θ*_ and selects a batch 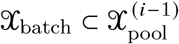. We then acquire the labels 𝒴_batch_ for 𝒳_batch_ (i.e., for our biological setting, we run the perturbation experiment). Finally, we update the labeled set 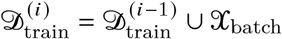 and pooled set 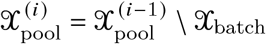. We proceed with the next round until a total of *R* rounds is reached.

Recently, the algorithmic framework by Holzmüller et al. [31] unified a large number of existing methods for this task. Their approach relies on reproducing kernel Hilbert spaces (RKHS), and computations on kernel matrices. Specifically, it consists of three steps. (1) *Base kernel calculation*. We construct a positive semi-definite kernel *k* : 𝒳× 𝒳 *→* ℝ to capture how the predictions from *f*_*θ*_ change with respect to 𝒳. A common choice is to build a finite-dimensional feature map *ϕ* : 𝒳 → ℝ^*d*^ with *k* (*x, x*′) = ⟨*ϕ*(*x*), *ϕ*(*x*′)⟩. Typical examples are the full gradient kernel, obtained for *ϕ*grad (*x*) = Δ _*θ*_ *f*_*θ*_ (*x*), as well as the last layer kernel 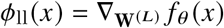, where W^(*L*)^ is denoted as the last layer parameter of the model *f*_*θ*_ . We note that because there are *L* gene expression levels to predict for each perturbation, we are interested in a multi-task prediction problem. Yet, we may still operate within this framework. Indeed, even though the gradient vector Δ_*θ*_ *f*_*θ*_ becomes a Jacobian matrix Jac *f*_*θ*_ (*θ*), we may just identify the matrix space to a finite-dimensional vector space. (2) *Kernel transformation*. While the base kernel defines the relation among inputs, it often requires an additional kernel transformation step for better performance, such as min-max normalization. (3) *Selection rule*. Lastly, given the transformed kernel, a selection method is invoked. The overall principle is to select informative and representative points that account for diversity. Since our proposed strategy modifies neither the kernel transformation nor the selection rules, we refer the readers to [31] for a more detailed review.

## 3 Method

### 3.1 Sequential design of Perturb-seq experiment

We now describe the unique challenges that may arise while designing an active learning strategy for the sequential design of Perturb-seq experiments.

#### Problem definition

We consider an initial Perturb-seq readout 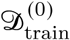 and a pool of unperturbed genes 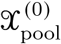 . In each round *i*, we train a perturbation prediction model *f*_*θ*_ using available data 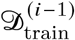 . Then, an active learning selection strategy *g* selects a batch 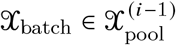 . We then conduct a wet-lab Perturb-seq experiment on these selected perturbations and obtain a batch of new readouts 𝒴_batch_, and proceed with the next round. The goal is find a selection strategy *g* that minimizes the model’s prediction error ℒ ^(*i*)^ on all the perturbations. For evaluation purposes, we evaluate the performance on a hold-out set of perturbations 𝒟_test_ at each round *i*.

#### Experimental setup

We focus on a CRISPRi Perturb-seq screen on cells from the K562 cell line undergoing 2,058 single-gene perturbations (essential genes as defined in [11]). We construct a benchmark to simulate the real-world active learning loop. First, we randomly select a hold out set of 205 perturbations for evaluation. This randomly selected hold out set gauges a model’s capacity to predict the entire perturbation space. Next, we set the number of rounds *R* = 5 and the number of perturbations that can be performed in each round *N*_batch_ = 100. For the sake of a fair comparison, we fix a random initial set of 100 perturbations for all methods. We measure the error at round *i* using the GEARS training loss at the hold out test set.

#### Economic analysis reveals active learning on a budget setting

Our problem is drastically different from the conventional active learning setting in several ways. First, previous works focus on single-output classification/regression tasks [32, 31], while the outcome of a Perturb-seq experiment is high-dimensional. This means that the predictive model may be harder to learn and thus may require more data.

Second, in a typical setting, the initial labeled set |𝒟_train_| is large enough for model training [32], followed by a large number of labels queried per round. However, this large number of labeled data is unattainable for Perturb-seq data, because each perturbation is associated with a high cost. Perturb-seq’s cost (currently dominated by the cost of scRNA-seq) for one perturbation can be estimated as the price of processing and sequencing a cell (varies across techniques, for droplet-based microfluidics, ∼0.5$ [33]) times the number of guides per perturbation (∼2) times the number of cells per guide (∼30) in addition to the pro-rated cost of labor, instruments, and quality control. Thus, a single perturbation is currently estimated to cost more than $30, making the number of perturbations intrinsically limited per round. Indeed, most Perturb-seq experiments reported to date are in the order of hundreds of perturbations [34, 8, 2, 10], largely driven by cost, as the experiment scales readily to genome-scale in the lab.

Third, previous works assume that many rounds of data acquisition can be performed. For example, the recent GeneDisco active learning challenge uses up to 40 rounds [35]. In contrast, each round of a Perturb-seq experiment is time-consuming. With some variation due to differences in the experimental platform, on average, each round of Perturb-seq takes at least a month (1 week for oligonucleotides synthesis, 1 week for library cloning, 1 week for titering, and 1 week for experiments). Thus, the number of rounds *R* should be small since the total time grows linearly with *R* for sequential Perturb-seq design. 40 rounds correspond to more than 3 years of implementation and is thus not realistic with current assay capabilities. This is why in our setting, we use *R* = 5.

Overall, we are operating in a different regime that we summarize as active learning on a budget [32]. We demonstrate below that this has significant impact on the design of the active learning strategy.

### 3.2 Data-driven motivation for incorporating prior knowledge

We hypothesize that the setting of active learning on a budget will affect the performance of any active learning strategy significantly because of the estimation of the kernel matrix, and therefore also the estimated relationships between perturbations, may be highly biased. We next present an analysis to support this hypothesis.

#### Testing alignment of kernels

We develop a simple test to gauge the quality of any kernel for downstream utilization in an active learning strategy. Since we have access to ground truth data (i.e., the outcome of all perturbations), we construct a ground truth kernel *k*_truth_ with *ϕ*_truth_(*x*) = y, where y denotes the experimental result. We expect the kernel matrix 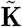 to reflect pairwise perturbation relationships, up to experimental noise. Therefore, the kernel matrix K derived from the predictive model *f*_*θ*_ should ideally be aligned as closely to the ground truth kernel as possible. To measure the alignment between the query kernel K and the ground truth kernel 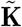, we use the kernel alignment score KA(K, 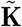) [36] defined as the cosine similarity between the two matrices (using the inner product canonically induced by Frobenius norm).

#### Poor alignment of the predictive model kernel when on a budget

We apply a baseline active learning algorithm for five rounds, where, at each round, we randomly query 100 new perturbations (random selection rule). At each of these five rounds, we also retrieve the perturbation prediction model, compute the kernel matrices and calculate the alignment score with ground truth. To make these kernel matrices comparable across rounds, we calculate them on the list of perturbations in the pool set of the last round. We observe that as the number of profiled perturbations increases, the model kernel alignment score also increases (Figure 1c). However, the alignment scores are low during the first few rounds, suggesting that the kernel matrix does not accurately represent the similarities between perturbations. This will lead to suboptimal selections since selection rules solely rely on the kernel to make selections.

**Figure 1:**
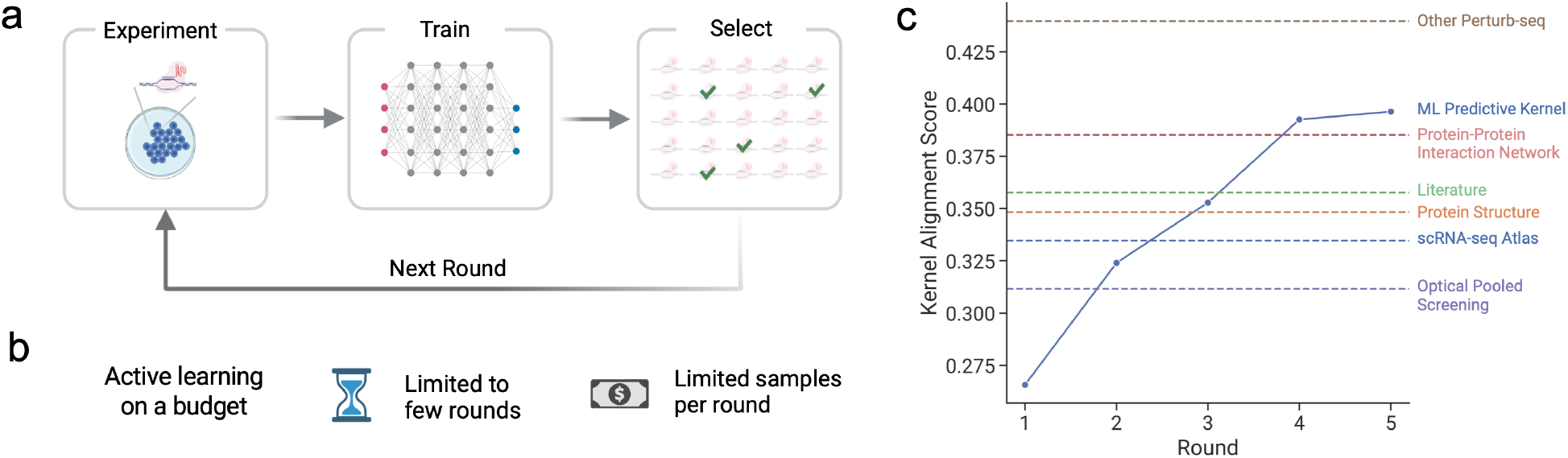
Sequential design of Perturb-seq experiments. **a**. Illustration of the iterative Perturb-seq procedure. In each round, a batch of perturbations is selected and the corresponding experiments are conducted. Then, a machine learning model is updated with these newly-profiled perturbations. An active learning strategy uses the model’s predictions to select the set of perturbations for the next round. Through this iteration, the goal is to reach high accuracy with a minimal number of experiments. **b**. Illustration of “active learning on a budget”. Active learning for Perturb-seq is highly restricted to much fewer profiled perturbations (i.e., labeled examples) compared to a conventional active learning setting. This motivates the development of a specialized method for this setting. **c**. Exploratory data analysis shows that the model kernel suffers from poor representation when few perturbations have been profiled (low budget). However, other data sources, described in Section 3.3, contain rich and complementary information that can be potentially transferred to the model kernel, motivating IterPert.

#### Prior knowledge contains auxiliary information of perturbation relationships

To tackle the insufficiency of the model kernel, we hypothesize that we can leverage abundant information about perturbation relationships stored in other sources of prior knowledge. To support this hypothesis, we collect a list of such sources and derive kernels that represent perturbation similarities (details about the sources and kernel derivations appear in Section 3.3). Using the same kernel alignment metric (Figure 1c), we observe that kernels derived from prior information have better alignment with the ground truth kernel compared to the model predictive kernel, especially in the first 2 rounds, suggesting rich information that could be complementary to the model kernel. This motivates us to design a method that integrates prior knowledge into active learning strategies.

### 3.3 IterPert: a multi-modal prior-guided active learning strategy

#### Overview

Motivated by the exploratory analysis in Section 3.2, we propose IterPert, an active learning strategy that incorporates diverse sources of prior knowledge to complement the model kernel when on a budget. The key step of our method consists in defining a kernel on each source of prior knowledge and combining those kernels with the model kernel to capture the relations between perturbations more accurately. This new prior-fused kernel is then followed by standard selection rules to form an active learning strategy.

#### Kernelized multi-modal prior information

Prior knowledge may come from diverse modalities, such as images, texts, and networks; therefore, how to employ these sources of prior knowledge for active learning is not straightforward. The information needed for active learning is not the raw prior knowledge, but the relations between the perturbed genes captured in the prior knowledge (e.g., using a kernel matrix). Thus, we propose to define a kernel *k* (*x, x*′) = ⟨*ϕ*_prior_ (*x*), *ϕ*_prior_ (*x*′)⟩ for each source of prior knowledge. Here, we introduce 6 distinct categories of prior knowledge, explain how to engineer a feature map *ϕ*, and provide insight on why each one should intuitively help map the perturbation space. The detailed preprocessing for each source can be found in Appendix A.

##### (1) Additional Perturb-seq data

Multiple Perturb-seq experiments have been conducted across several cell contexts [34]. Perturb-seq data from other cell contexts or experiments contain useful prior information since certain relations between perturbations might be either context-agnostic or at least transferable to the cell context of interest. For each perturbation *x, ϕ* (*x*) is defined as the mean of pseudo-bulk expression change from the non-targeting control cells from the Perturb-seq readouts.

##### (2) Optical pooled screens (OPS)

OPS [24, 25] data consists of cell morphological images associated with a genetic perturbation in each individual cell in a pool. Intuitively, perturbations that elicit similar morphological phenotypes could also have similar expression phenotypes. For each perturbation *x, ϕ x* is the imaging features from CellProfiler [37], an image processing software that extracts morphology profiles.

##### (3) scRNA-seq atlas

Genes that are co-expressed together likely belong to similar pathways, and perturbations in the same pathway tend to have similar expression effects. Thus, gene co-expression data could be useful for the prediction task. For each perturbation *x, ϕ x* is the list of normalized gene expression measurements for gene *x* across a collection of scRNA-seq experiments [38]. The kernel matrix derived from this feature map corresponds to the co-expression matrix.

##### (4) Protein structures

If the proteins encoded by the perturbed genes have similar structures, they are more likely to have similar functions, and similar perturbation outcomes [26]. For each perturbation *x*, we obtain its protein coding sequence, and then feed it into a recent protein language model (15B ESM model [27]) to obtain structural features *ϕ*(*x)* .

##### (5) Protein-protein interaction network (PPI)

A PPI network connects proteins that physically interact with each other [39]. Intuitively, a physical interaction between two proteins suggest that they might participate in a shared biomolecular pathway or complex. Thus, perturbations of genes coding for physically interacting proteins might lead to similar effects [40]. For each perturbation *x, ϕ x* is a node embedding of *x* in the PPI network, such as node2vec [41].

##### (6) Literature

Perturbations that are mentioned in similar contexts in the literature are more likely to have similar functions and phenotypes. To encode this, for each perturbation *x, ϕ x*, we feed the corresponding gene name to a recent large language model that is fine-tuned on biological literature (*e*.*g*. BioGPT [42]) and use the text embedding as the feature map.

#### Kernel fusion

The kernelization step enables integration across kernels with diverse modalities, since it converts different modalities into one — a kernel matrix of the same size. Now, we study how to fuse all the prior kernels with the model kernel. Given the set of prior kernels {*k*_1_, …, K_*m*_} and their kernel matrices {K_1_, …, K_*m*_}, we update the model kernel matrix K at each round to obtain the kernel matrix 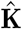 for our active learning procedure as follows:

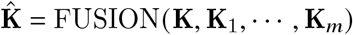

Since the different prior kernels have different feature map dimensions, and the kernel corresponds to taking the dot product between feature maps, the scale differs significantly across kernels. Thus, to avoid one kernel with a large scale overriding the others, we apply a min-max scale normalization to each kernel.

We experiment with multiple strategies for the FUSION operator, including element-wise operators, such as mean, max, and product, and adaptive kernel aggregation methods, such as the kernel alignment weighted operator and the kernel regression operator. A discussion and performance study of these fusion operator appears in Appendix B. Interestingly, the mean operator 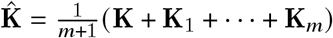 has the best empirical performance. Additionally, this approach has the theoretical advantage of guaranteeing that the fused kernel is positive and semi-definite (PSD), which is a required property for several downstream selection rules [31]. Note that the mean operator also has an interpretation in the feature maps space, where it is equivalent to the concatenation of all the feature maps.

#### Selection rule

IterPert only modifies the base kernel and is agnostic to the selection rule. For the sake of simplicity, in our experiments, we apply IterPert with only one popular rule called greedy distance maximization. This method greedily select points with maximum distance to all previously selected points [43]. Particularly, given the prior-fused kernel 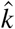, for a perturbation *i* in the pool set and any point *j* in the selected set, it first calculates the distance 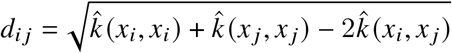. This is equivalent to taking the squared distance in the feature map space. Next, it selects point *i*\* greedily as

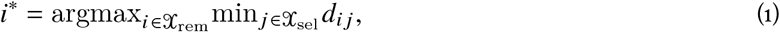

where 𝒳_sel_ is the union of the training set and the points already selected, and 𝒳_rem_ is the pool set excluding the already selected points.

## 4 Experiment

We conduct experiments to demonstrate IterPert’s advantage over state-of-the-art active learning strategies in efficiently designing Perturb-seq experiments. We also conduct systematic ablation studies to delineate the contribution of each prior information source. We evaluate the performance of our benchmarked methods in various settings, including an extension to a larger pool size by leveraging a genome-scale Perturb-seq screen and also accounting for batch effects across rounds.

### Benchmarking state-of-the-art active learning methods

We first benchmark the set of active learning methods (Figure 3b) available from the open-source repository released by Holzmüller et al. [31]. We observed that all active learning methods have better performance than uniform/random sampling. The best-performing method was TypiClust [32], which is a recent active learning on a budget method that prioritizes typical examples instead of uncertain examples and shows significant improvement over random selection. This corroborates our hypothesis that the problem of sequential Perturb-seq experimental design corresponds to the setting of active learning on a budget.

**Figure 2:**
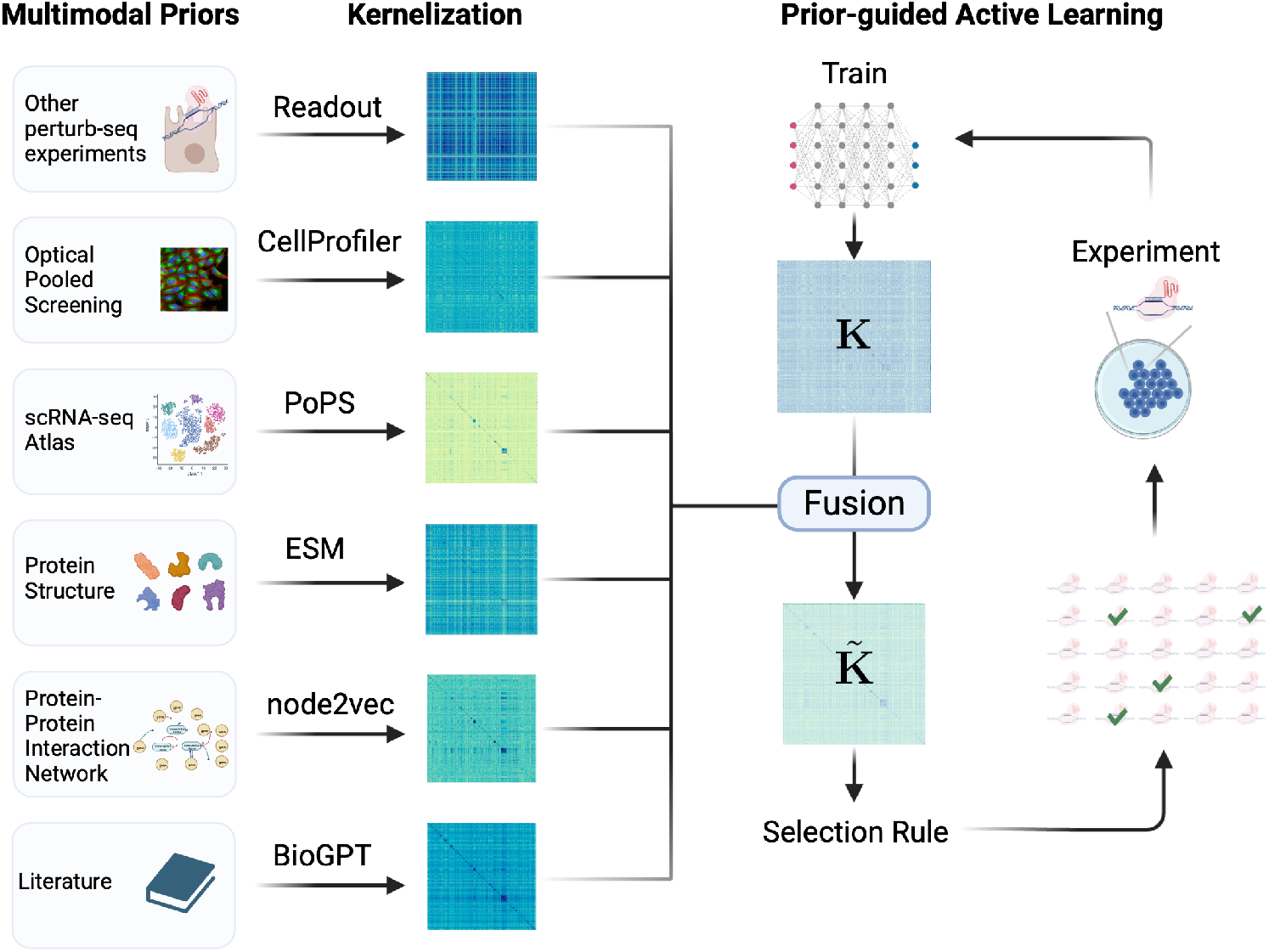
Illustration of IterPert. Driven by the exploratory data analysis in Section 3.2, we introduce IterPert, an active learning selection approach that integrates a wide range of multi-modal prior knowledge to tackle the problem of active learning on a budget for Perturb-seq data. Our primary technique involves enhancing the model kernel when faced with budget constraints. We achieve this by transforming each source of multi-modal prior knowledge into a reproducing kernel Hilbert space through diverse featurization methods for each modality, explained in Section 3.3 These kernels are then fused to refine the model kernel, ensuring a more precise characterization of perturbation relations. We then apply standard selection rules to this enhanced kernel, giving rise to IterPert.

**Figure 3:**
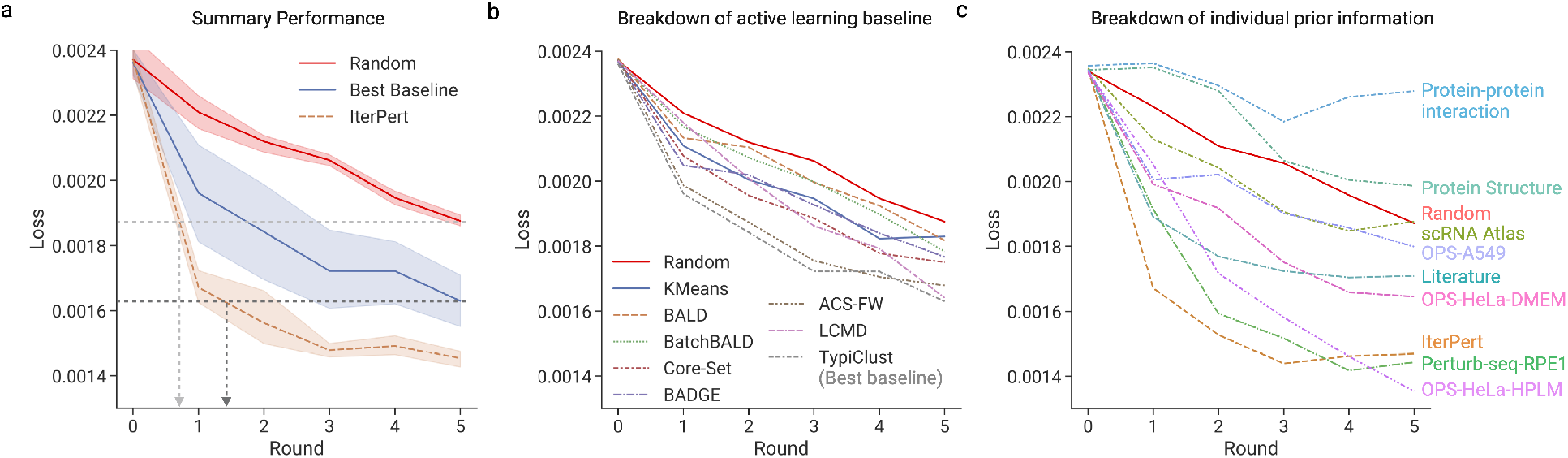
**a**. IterPert achieves significant speedup of model learning compared to the best baseline and random selection. Focal loss (training objective of the base model, y axis) across active learning rounds (x axis). We conduct 10 random runs where the solid line is the average and the error bar is the 95% confidence interval of the mean. **b**. Detailed breakdown of state-of-the-art active learning baselines. The best baseline is TypiClust [32]. Plot as in panel a, with the solid line denoting the average across 10 runs. **c**. Detailed breakdown of individual prior-augmented active learning. The solid line is the average across 5 runs. Error bars are not visualized in panels b and c for visual clarity and can be found in Appendix C.

### IterPert achieves significant improvement over the best baseline

We report the performance of IterPert against the best active learning baseline and random sampling in Figure 3a. Importantly, IterPert uses roughly one round to reach the same accuracy as five rounds of uniform sampling, reflecting a greater than 5-fold speedup. Similarly, IterPert uses roughly 1.5 rounds (through linear extrapolation) to reach the same accuracy as five rounds of uniform sampling, reflecting a more than 3-fold speedup. We also observe similar improvements in other biologically meaningful metrics, such as the mean squared error (MSE) of predicted expression profiles calculated on the top 20 differentially expressed genes in each perturbation (Figure 4a) and the Pearson correlation coefficient over changes in gene expression (Figure 4b). This showcases that IterPert is an efficient method for designing Perturb-seq experiments. Moreover, the first round had the steepest increase of accuracy for IterPert, confirming our data analysis in Section 3.2 on the usefulness of prior knowledge.

**Figure 4:**
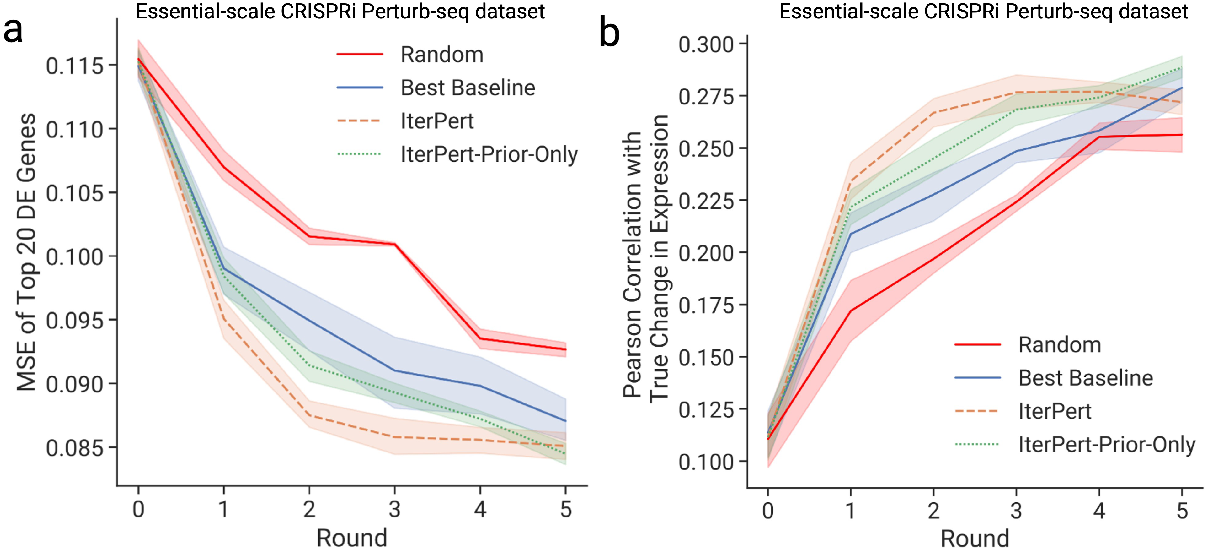
Biologically meaningful metrics as used in [12]. Metrics (y axis) across active learning rounds (x axis). Each method is averaged across 10 runs and error bar is the 95% CI of the mean. **a**. MSE of top 20 differentially expressed genes per perturbation. **b**. Pearson correlation coefficient between the predicted and true expression changes (centered on non-targeting controls). IterPert-Prior-Only is an ablation of IterPert where we remove the model kernel. Best baseline is TypiClust [32].

### Dissecting IterPert performance across multi-modal priors

To further understand the origin of the performance improvement of IterPert, we conduct several ablations. First, we report performance when using a single prior kernel, so that we may understand which prior source contributes most to the performance of the method (Figure 3c). Aggregation of all priors outperformed any individual prior alone. This showcases synergies across the diverse sources of prior knowledge. Comparing across priors, the best-performing prior is the Perturb-seq data in RPE1 cells, highlighting that there is transferable information across Perturb-seq experiments, even from different cell contexts. Optical pooled screens were also strongly informative, demonstrating that cell morphology carries shared information with Perturb-seq outcomes. Notably, different cell contexts and treatment/phenotypes of OPS lead to different improvement levels. The HeLa cell line seems to have a larger contribution to model performance increase than an OPS in the A549 cell line. Other prior knowledge sources, such as literature and a scRNA-seq atlas, also show an improvement, while PPI has limited contribution, maybe due to noise. Overall, perturbation-specific priors have richer signals compared to general gene-based priors. We also conduct an ablation where we remove the model kernel (Figure 4a,b). We observe a performance degradation, highlighting the synergy between prior knowledge and the model kernel.

### Extension to genome-scale experiment

The pool set of the essential genes in K562 dataset is relatively small (<2,000 perturbations). For many real-world applications of Perturb-seq, one may want to select from a larger pool set size, for example, in genome-scale screens or in combinatorial screens. To gauge the improvement in larger setups, we conducted another experiment by leveraging the genome-scale K562 CRISPRi perturb-seq screen from [11]. This dataset has 9,748 single-gene perturbations and thus corresponds to a much larger pool of possible perturbations. We set *N*_batch_ = 300 and performed *R* = 3 rounds in total. We report the performance in Figure 5. We find that IterPert consistently displays a significant efficiency improvement over both the random and best active learning baselines, especially in the first round.

**Figure 5:**
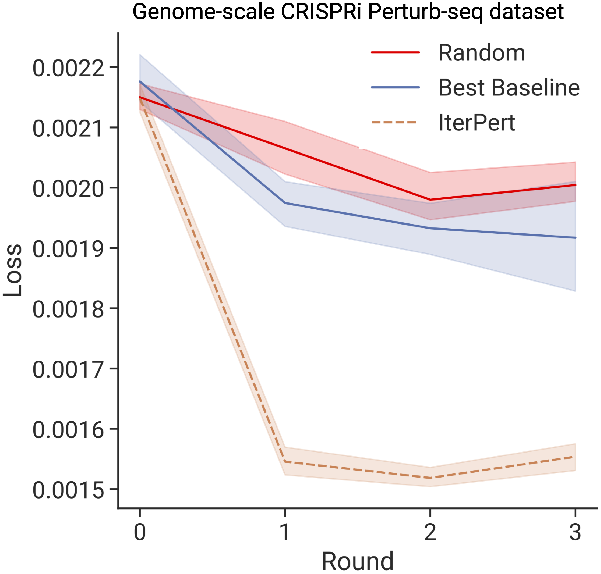
Performance for genome-scale K562 CRISPRi screen. Focal loss (y axis) across active learning rounds (x axis). Solid line: mean over 10 runs, error bar: 95% CI of the mean. Best baseline is TypiClust [32]. Results for other baselines can be found in Appendix D.

### Accounting for batch effects across rounds

One important consideration when developing an active learning strategy for Perturb-seq data is that there are batch effects across rounds (Figure 6a), which could bias the predictive model and selection strategy. To evaluate the our method’s robustness to this, we simulate batch effects by leveraging the batch information in the dataset, which consists of 48 batches (lanes) (Figure 6b). In particular, we restrict the cells for 𝒳_batch_ in each round to come from different batches (8 batches for each round), to ensure that the model experiences some batch effects. We conduct the same experiment and report the performance in Figure 6c. We observe that the absolute value of the loss is worse than in the previously explored settings without batch effects. This may mainly be due to the fact that 6 times fewer cells are available for training. In this challenging setting, we still observe that IterPert has a more efficient selection strategy compared to the best baseline and uniform sampling.

**Figure 6:**
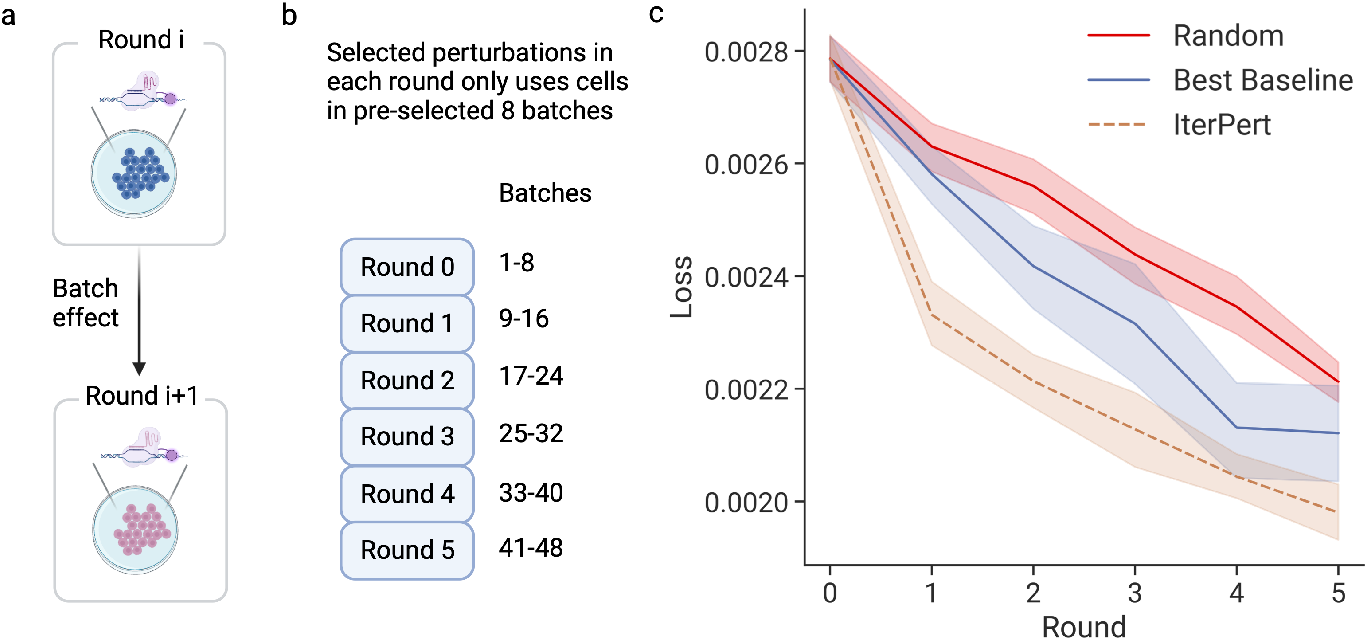
**a**. Batch effects exist across active learning rounds, which could bias model training and selection. **b**. Illustration of simulation evaluation settings where we restrict the cells from the selected perturbations in each round to certain batches (lanes) such that different rounds use cells from different batches. **c**. Active learning performance in the batch effect setting. Each method is averaged across 10 runs and error bar is 95% CI. Best baseline is TypiClust [32].

## 5 Related works

### Active learning

We highlight recent advancements that we consider as baselines and refer the readers to surveys [15, 31, 44] for a more comprehensive overview. BALD [45] selects instances where the model’s predictions exhibit the most disagreement across possible parameter configurations, focusing on uncertainty. Batch-BALD [46] is an extension of BALD and it selects batches of data points to maximize joint information and reduce redundancy in batch selection. Core-Set [43] identifies a subset of data that summarizes the entire dataset, aiming for comparable performance with fewer training examples. BADGE [47] chooses data points based on diverse gradient embeddings, capturing instances that offer varied learning experiences. ACS-FW [48] uses the Frank-Wolfe optimization algorithm to select instances from the pool set whose conic combinations best represent the entire set to promote representativeness. LCMD [31] first finds the largest cluster for representativeness and then enforces diversity by picking the maximum distance point within this cluster.

### Active learning on a budget

Active learning on a budget has been studied in [21, 22, 23]. They showed that in this setting, random selection outperforms most deep active learning strategies. This phenomenon is often explained by the poor ability of neural models to capture uncertainty on a small budget. The recently proposed method TypiClust [32] prioritizes typical examples instead of uncertain examples and shows significant improvement over random selection. We consider it as our baseline. Note that with IterPert, we do not propose a new selection rule but instead use prior information to adjust the estimation of the perturbation space. We show that IterPert has significant improvement over TypiClust, but we leave the problem of integrating IterPert with TypiClust as future work.

### Perturbation prediction models

CellOracle[29] relies on gene regulatory network inference and conducts linear network propagation of perturbation signals to make predictions. CPA[30] uses a non-linear compositional autoencoder to predict effects but it is restricted to predicting seen perturbations. GEARS [12] is a deep learning model customized for perturbation prediction. It is based on a GNN perturbation and cell encoder with a deep composition layer that simulates multi-gene perturbations on cells, and it features a loss function focusing on differentially expressed genes. Recently, single-cell foundation models have gained popularity and claim to excel at perturbation outcome prediction. Notably, scGPT [13] uses a generative pre-training objective over a massive scRNA-seq atlas and is finetuned on perturbation prediction tasks. However, it requires the perturbed genes to be detected in the scRNA-seq experiment, which is not the case for many perturbations in our data set. Although we use GEARS in this work, the approach is general and applicable to other models.

### Active learning for genomics experimental design

Sequential optimal design is increasingly popular in high throughput genomics assays. The main task is to identify genes that maximize an endpoint such as cell proliferation [49, 50]. Note that this setting is highly different from ours, because there, the goal is to identify a data point in the data distribution with the highest response (Bayesian optimization). In contrast, we are interested in selecting points that enable a machine learning model to reduce the overall loss across the data distribution (active learning). The more related work GeneDisco [35] is a benchmark for the sequential design of genetic perturbation experiments, proposing both Bayesian optimization and active learning tasks. The key difference in our work is that we focus on active learning for expensive Perturb-seq, where the response is high-dimensional expression profiles, while GeneDisco focuses on functional genomics CRISPR assays with a single scalar readout. This leads to different base prediction models and a different active learning setting than the one discussed in Section 3.1. Also note that we have included the active learning methods benchmarked in GeneDisco (BADGE, KMeans, BALD) in our baselines, and in this study, our proposed method IterPert has significantly better performance.

## 6 Discussion

We introduced an iterative Perturb-seq procedure for efficient design of perturbation experiments. We highlighted the challenges of active learning on a budget constraints and evaluated current active learning techniques. Motivated by an initial data analysis, we presented IterPert, a new active learning strategy that incorporates multi-modal priors, achieving over three times the speed of the best baseline.

While IterPert shows promise in designing efficient Perturb-seq experiments, it still faces limitations, and further work is necessary for its practical implementation. For instance, while we strive to simulate a realistic setting *in-silico*, several points of divergence could occur in practice. One such divergence is experimental batch effects, which could be more significant than those considered in our setting. Moreover, while our method is very useful for mapping genome-scale single-gene perturbations, further work is needed to extend this approach to multi-gene (combinatorial) perturbations that are currently intractable to experimentally interrogate in an exhaustive way. Extending the framework to multi-gene perturbations requires higher-order kernels or the use of tensor product spaces, which presents an interesting methodological challenge that we leave for future work. Similarly, extensions to chemical perturbations or optical readouts are also exciting future avenues. More specific to our prior-guided strategy, while our empirical study finds that mean fusion works the best, it is not context-specific. Ideally, different combinations of prior information could be automatically picked in different cell contexts. Lastly, with the increasing interest in models to predict the outcome of perturbations, we expect more base prediction models to become available. While our proposed active learning strategy is compatible with any of these, future work remains to investigate IterPert performance with these methods.

Overall, we believe that the sequential design of Perturb-seq could drastically reduce the experimental cost of understanding a complex space of perturbations, thanks to its sample efficiency, and could help answer central biological questions, such as the effect of multi-gene perturbations.

## Acknowledgements and Funding Information

We thank Xinming Tu, Jerry Wang, and Rebecca Boiarsky for feedback throughout the duration of this project that greatly improved this work. We warmly thank Jure Leskovec for valuable discussions and feedback for improving the manuscript. We also thank members of the Regev Lab and the Biological Research | AI development (BRAID) department at Genentech for providing constructive feedback on earlier versions of the results presented in this work.

Romain Lopez, Jan-Christian Hütter, Takamasa Kudo, Antonio Rios, and Aviv Regev are employees of Genentech, and may have equity in Roche. Aviv Regev is a co-founder and equity holder of Celsius Therapeutics and an equity holder in Immunitas. She was an SAB member of Thermo Fisher Scientific, Syros Pharmaceuticals, Neogene Therapeutics, and Asimov until July 31st, 2020.

## Code Availability

The raw source code is available at https://github.com/Genentech/iterative-perturb-seq and is released under the Apache 2.0 license. The notebooks to reproduce each figure are provided in https://github.com/Genentech/iterative-perturb-seq/tree/master/reproduce_repo. The python package is available at iterpert. We implemented the source code in PyTorch. The base machine learning model is adapted from https://github.com/snap-stanford/GEARS. The active learning strategy framework is adapted from https://github.com/dholzmueller/bmdal_reg.

## Appendix

In Appendix A, we describe pre-processing steps for each prior source we leverage. In Appendix B, we describe different fusion operators to fuse across prior kernels and report their empirical performance. In Appendix C, we further provide plots of the main experiments that include all error bars since they are withheld for the sake of visibility in the main text. In Appendix D, we discuss the experiments on the genome-scale Perturb-seq data and report the obtained performance metrics.

### A Data processing on multi-modal priors

1. Additional Perturb-seq data: we use the essential-wide RPE1 cell line CRISPRi dataset from the same paper [11] as the K562 dataset. Particularly, for each perturbation, we obtain the NTC centered pseudobulk expression profile and use that as the feature embedding. For genome-scale experiment, since we do not have another cell line with genome-scale perturbations, we remove this prior source.
2. Optical pooled screens: [25] conducts a genome-wide optical pooled screen and calculated CellProfiler features for each perturbation. We retrieve each perturbation embedding from https://github.com/broadinstitute/2022_PERISCOPE#downloading-profiles. Notably, we use the median aggregation version. For A549, we used 20200805_A549_WG_Screen_guide_normalized_median_merged_ ALLBATCHES_ALLWELLS.csv.gz. For HeLa, we used both DMEM 20210422_6W_CP257_ guide_normalized_median_merged_ALLBATCHES DMEM ALLWELLS.csv and HPLM 20210422_6W_CP257_guide_normalized_median_merged_ALLBATCHES HPLM ALLWELLS.csv.
3. scRNA-seq atlas: we used processed scRNA profiles aggregated from multiple scRNA-seq experiments in [38] (https://github.com/FinucaneLab/pops).
4. Protein structures: we retrieve the protein coding sequence of the corresponding gene perturbation from uniprot and then feed each into ESM-2 15 billion parameter model(https://huggingface.co/facebook/esm2_t48_15B_UR50D) and the output [CLS] token embedding is used as the protein embedding.
5. Protein-protein interaction network: we used the PPI knowledge network from https://arxiv.org/abs/2306.04766, and apply node2vec (https://github.com/eliorc/node2vec) to obtain each gene embedding.
6. Literature: we feed the gene name of each perturbation into BioGPT (https://huggingface.co/microsoft/BioGPT-Large) and we use the [CLS] token embedding as the gene embedding.

#### B Fusion operator

We experiment with multiple strategies for the FUSION operator, including element-wise operators:

1. Mean operator: 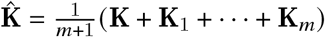
2. Max operator: 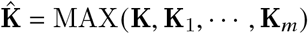
3. Product operator: 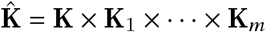

We also experiment with adaptive kernel aggregation methods. Given the subset of kernel matrix with ground truth at round *i* called 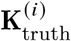, we can estimate the kernel alignment scores 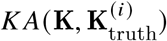, where *K A* [36] is defined as the cosine similarity between the two matrices (using the inner product canonically induced by Frobenius norm). The *kernel alignment weighted operator* is then defined as

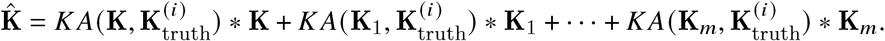

Another learnable operator is to estimate the weights *α, α*_1_,, *α*_*m*_ by solving a linear regression problem to fit the ground truth sub-kernel from prior sub-kernel using validation dataset at each round *i*:

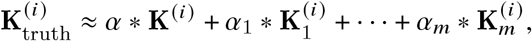

and then use the weights to update the entire kernel

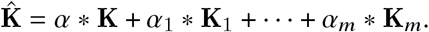

We report performance comparisons of these different operators in Figure 7. We observe that the mean operator has the best empirical performance. We hypothesize that the reason for this that while the learnable operators can capture context-specific relations among the kernels, their estimation is biased due to the limited size of available data for each round. We also experimented with non-linear integration of kernels, but they easily led to overfitting. In the end, we adopt the mean fusion operator.

**Figure 7:**
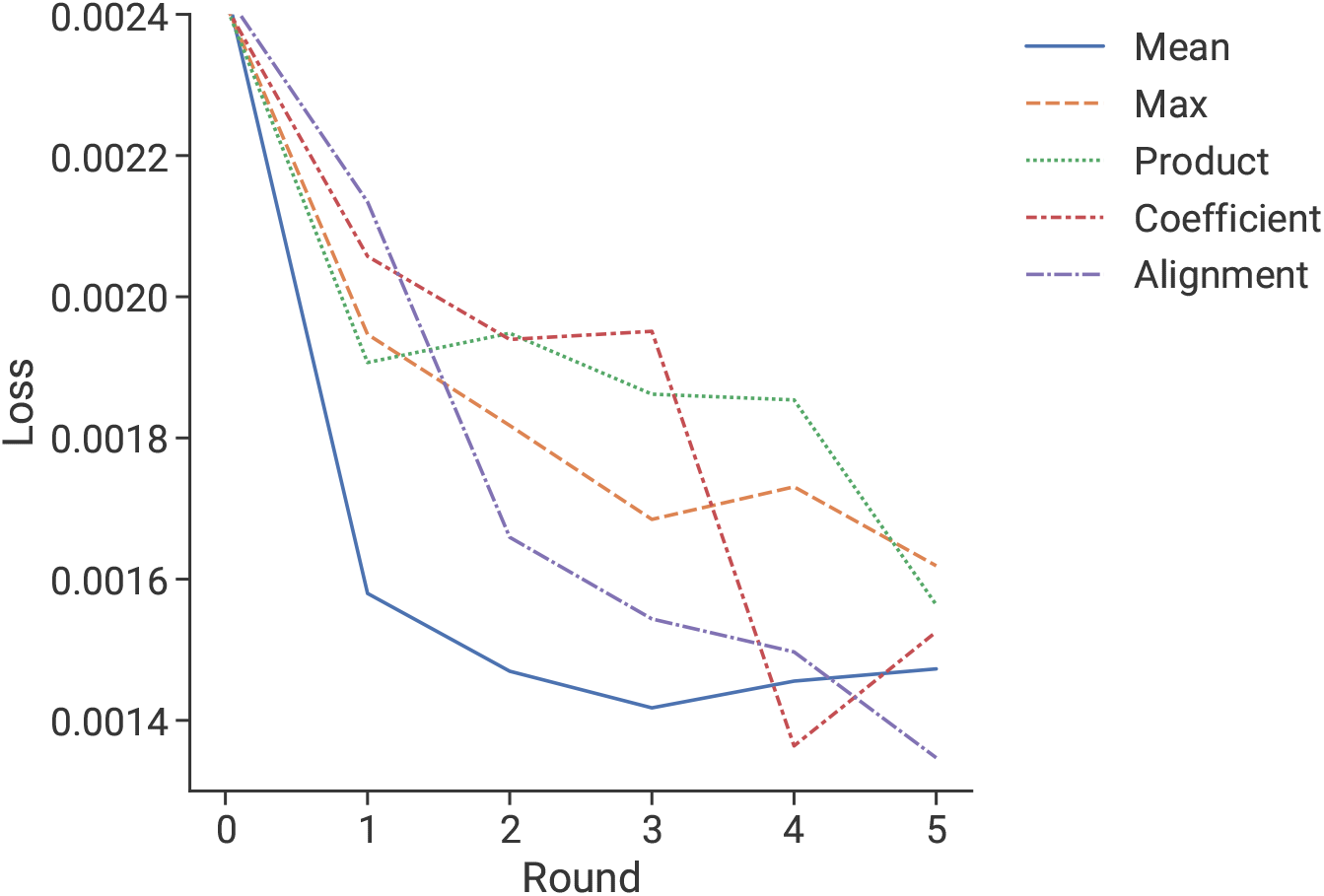
Performance comparison on a seed different from the main experiments across different fusion operators. Mean operator has the best empirical performance.

#### C Error bars for baselines

Error bars are omitted in Figures 3b and 3c in the main paper to make the plots easier to read. We here report the error bar for Figure 3b (breakdown of active learning baselines) in Figure 8 and the error bar for Figure 3c (breakdown of individual prior information) in Figure 9.

**Figure 8:**
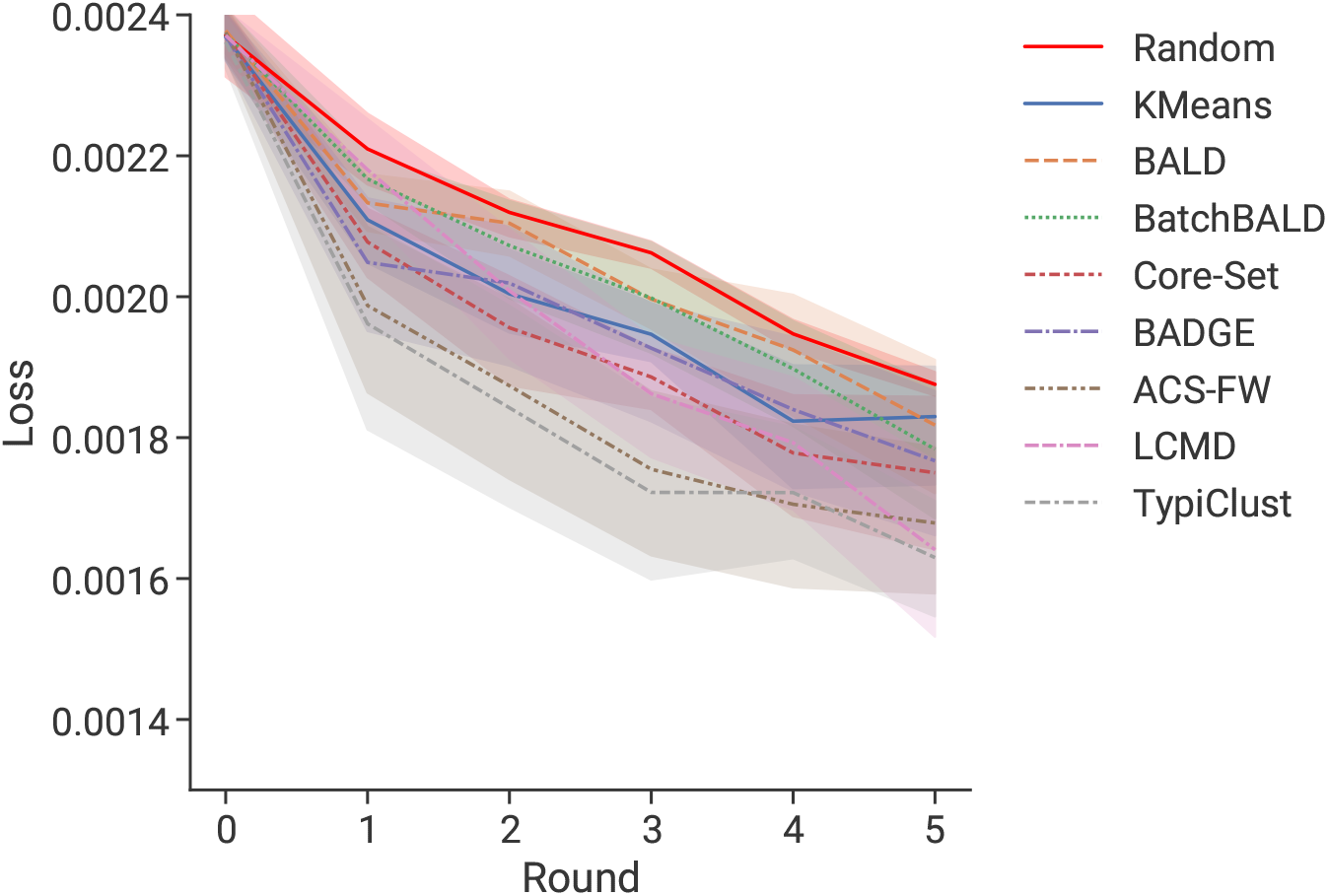
Performance comparison across different baselines with error bar corresponding to 95% confidence interval.

**Figure 9:**
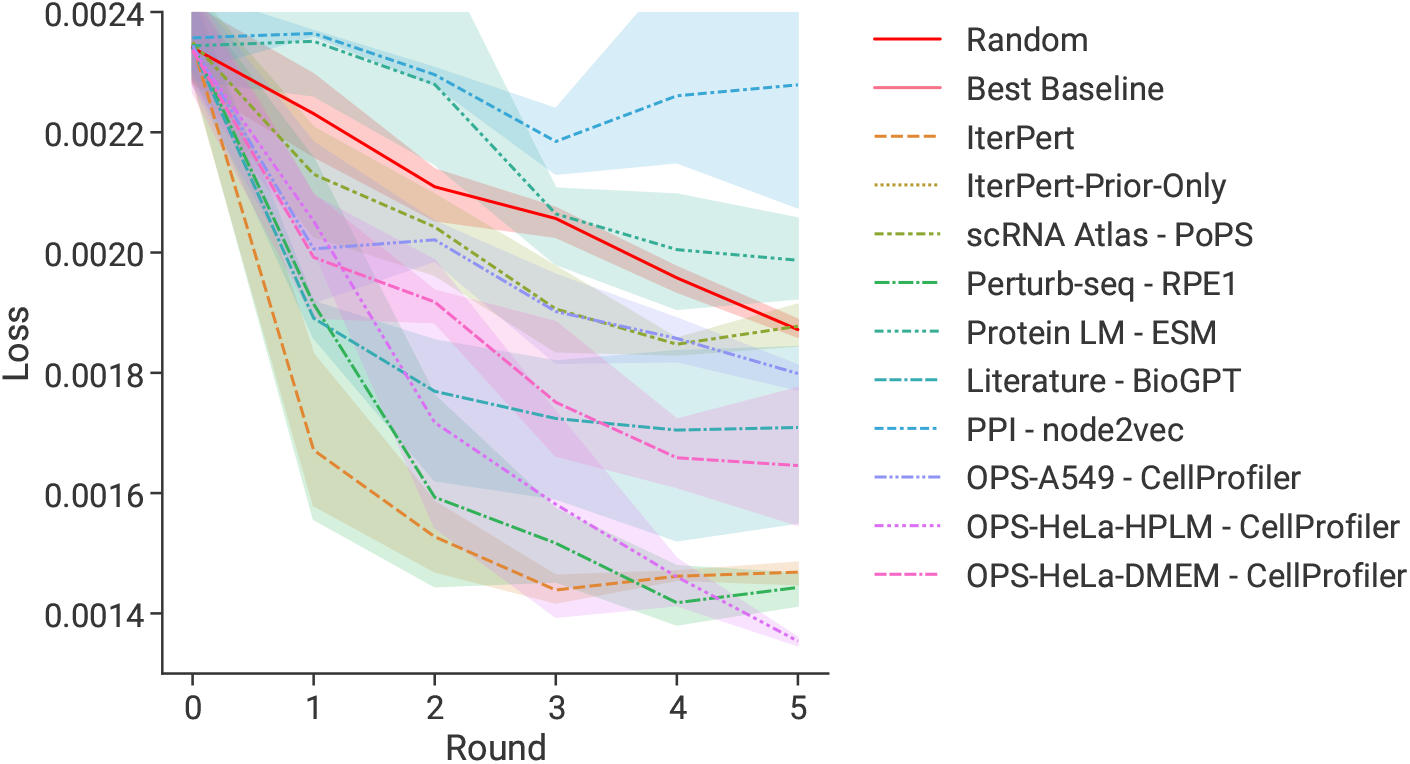
Performance comparison across different prior information with error bar corresponding to 95% confidence interval.

#### D Baseline performance for genome-scale perturbation screen

In Figure 10, we report the performance of all the baseline state-of-the-art active learning strategies on the genome-scale perturbation screen that were omitted in Figure 5.

**Figure 10:**
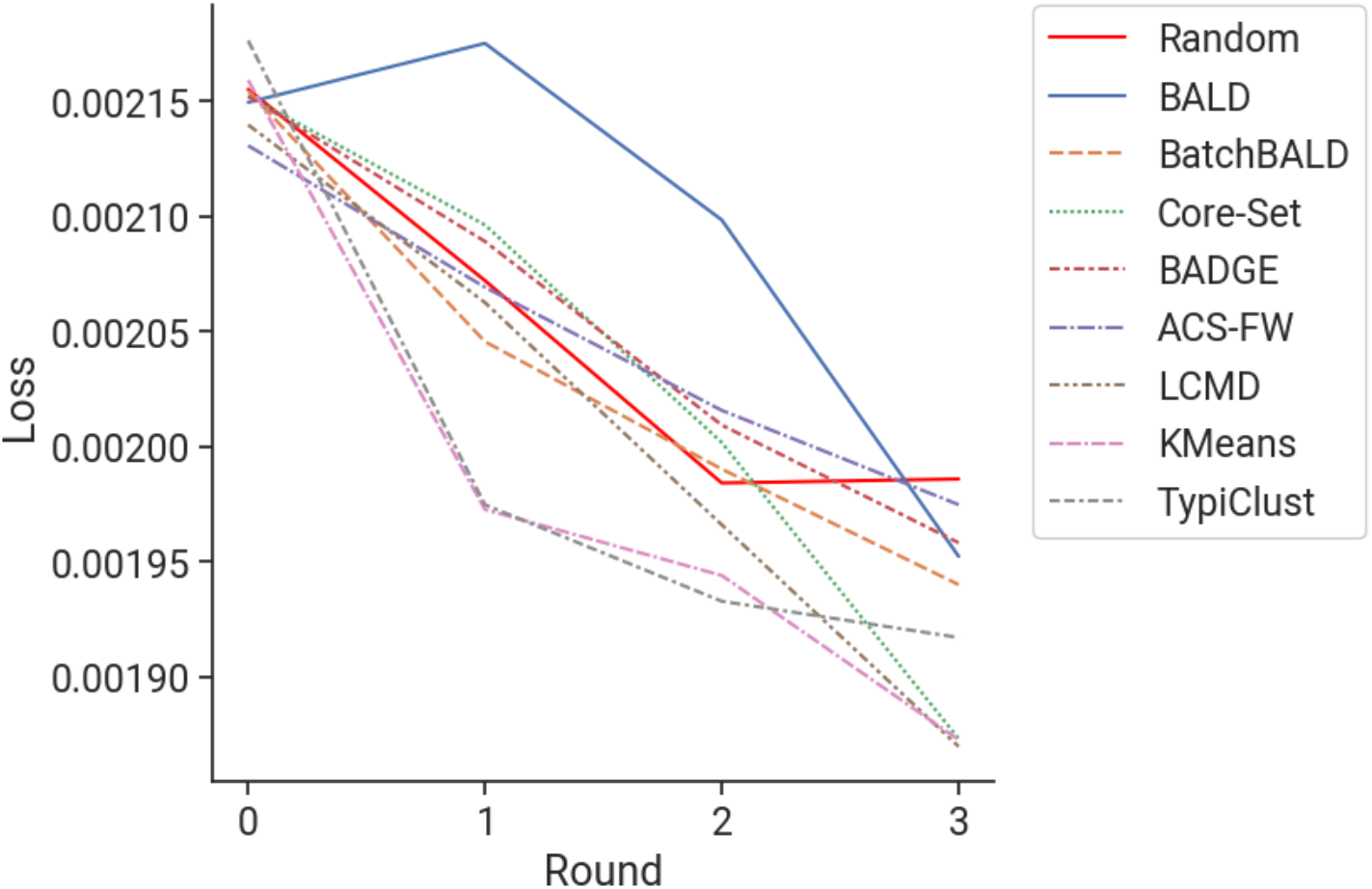
Performance comparison across baseline methods for genome-scale K562 screens with error bar corresponding to 95% confidence interval.

